# RamanMAE: Masked Autoencoders Enable Efficient Molecular Imaging by Learning Biologically Meaningful Spectral Representations

**DOI:** 10.1101/2025.05.18.654618

**Authors:** Santosh Kumar Paidi, Parul Maheshwari

**Affiliations:** Genentech; PayPal

## Abstract

Traditional histopathological analysis of cells and tissue relies on morphological features from stained biopsy samples, which fail to leverage the wealth of chemical information about the underlying pathological states. Raman spectroscopy, a form of vibrational spectroscopy, uses light scattering to capture chemical information about the biological specimen. However, advancements in Raman spectroscopy are hindered by the method’s intrinsic low throughput and the difficulty in deriving meaningful insights from the high-dimensional noisy datasets. In this paper, we propose RamanMAE, a spectral language model using masked autoencoders on large Raman spectral datasets from biological applications that can be used for spectral processing in applications with limited data. We achieved excellent reconstruction of masked patches of the spectra. We learned meaningful latent representations of the spectra that capture biological compositional information and serve as a low-dimensional feature space for building downstream machine learning methods. We showed that the decoder serves as an effective smoothing technique to reduce noise in the spectra and allow better localization and visualization of biological features in the spectral maps. We also demonstrated the transferability of the representations learned on one dataset to a different biological application.

## Introduction

The transformer-based masked autoencoders, a form of self-supervised learning, are scalable learners of features relevant to several domains (1–5). The initial success of BERT in natural language processing paved the way for the development of VIT-MAE by leveraging the vision transformer model (2, 6). It spurred numerous applications in computer vision and related application areas such as optical microscopy and perception (7–9). Motivated by the success of these efforts, we propose RamanMAE - a masked autoen-coder approach for learning representations of Raman spectroscopy data. Raman spectroscopy is a form of vibrational spectroscopy that probes the inelastic scattering of light to study chemical and biological systems (10, 11). Raman spectra capture rich molecular information about the underlying samples. With the advent of powerful lasers and sensitive detectors, the field of biomedical Raman spectroscopy has seen a surge in applications in biology and medicine in recent decades, ranging from clinical diagnostics and pre-clinical biological studies to analysis of biofluids for diagnostic and forensic purposes with unprecedented sensitivity and molecular specificity (10–12). However, since only one in a million photons are Raman scattered, the method traditionally lags behind other optical methods, such as multi-photon microscopy, in terms of throughput and typically requires large acquisition times to obtain spectra of acceptable quality. A variety of traditional machine learning methods, including linear methods such as partial least squares and non-linear methods such as random forests and support vector machines, are routinely used for inferring chemical and biological attributes of biomaterials from Raman spectra (13–15). These methods, while suitable for low-data applications, often require additional dimensionality reduction to prevent overfitting and need redevelopment for different applications.

The adoption of deep neural network architectures in Raman spectroscopy has been hindered by the lack of sufficiently large spectral datasets (16). Some researchers have used 1D convolutional neural networks for pre-processing Raman spectra, but the applicability of such methods for biological datasets is limited due to the use of synthetic datasets to train such networks (17–19). Recently, a few groups have obtained large spectral datasets for training application-specific deep neural network models and explored their transferability to new applications (20–22). Horgan et al. developed DeepR framework that leverages 1D ResUNet to produce high-quality output Raman spectra from low signal-to-noise ratio (SNR) input spectra that are strongly correlated with the target (ground truth) high SNR spectra (20). The authors did not assess the utility of the learned representations for downstream classification or regression tasks. Wu et al. designed a data augmentation module that employs a generative adversarial network (GAN) to generate synthetic Raman data resembling the training data classes, but the applicability of such networks trained on an application with unknown distribution is not fully studied (23). Some groups have also demonstrated the use of variational autoencoders with convolutional frameworks for clustering and classification in the latent feature space (24, 25). These methods are developed for specific applications and their transferability is not well explored.

In this paper, we explored the possibility of directly leveraging the VIT-MAE architecture with some key modifications to learn biologically relevant representations of Raman spectra for smoothing and downstream applications. Masked autoencoders are particularly attractive for Raman spectroscopy because (1) they learn low-dimensional representations (embeddings) of the sparse correlated datasets, (2) the learned embeddings are useful for downstream applications like classification and regression, and (3) the models generalize well to different applications beyond their initial training context.

Raman spectra are one-dimensional data but share similarities with image data due to the high correlation and high dimensionality of the spectral features. Therefore, by leveraging large Raman spectral datasets from published biological studies, we trained RamanMAE – a masked autoencoder model derived from the VITMAE model with some domain-specific modifications for domain adaptation **Figure 1**. Furthermore, the suitability of VITMAE for adaptation to Raman spectroscopy is justified by the serial patch encoding proposed in VIT (and by extension in VITMAE) that can be translated to serial patches of Raman spectra, which typically capture Raman peaks related to vibrational modes (2, 26). The use of flattened patches for serial positional encoding in VIT makes it suitable for 1D Raman spectra by reshaping the spectra appropriately. We show excellent reconstruction of Raman spectra and observe the inherent spectral smoothing of the reconstructed spectra, consistent with the smoothing behavior observed in masked autoencoders due to suppression of noise and improvement of signal to noise (SNR). We inspect the acquired low-dimensional embeddings and show that they capture biologically relevant features of the dataset by identifying key structural features of single biological cells. Next, by training the model on a challenging metastatic cancer cell Raman dataset, we demonstrate the utility of learned representations in predicting the metastatic phenotype at single cancer cell resolution. Finally, we explore the transferability across scale and application domain by training RamanMAE on one application and testing it for a different downstream task. These results demonstrate that masked autoencoders are scalable learners of biologically relevant features of Raman spectra in low-dimensional space and serve as inherent smoothing functions. We envision that with emerging cross-institutional data-collection efforts in the field of biological Raman spectroscopy, RamanMAE and masked autoencoder-based architectures in general are going to be valuable tools in the arsenal of Raman data analysis.

**Figure 1.**
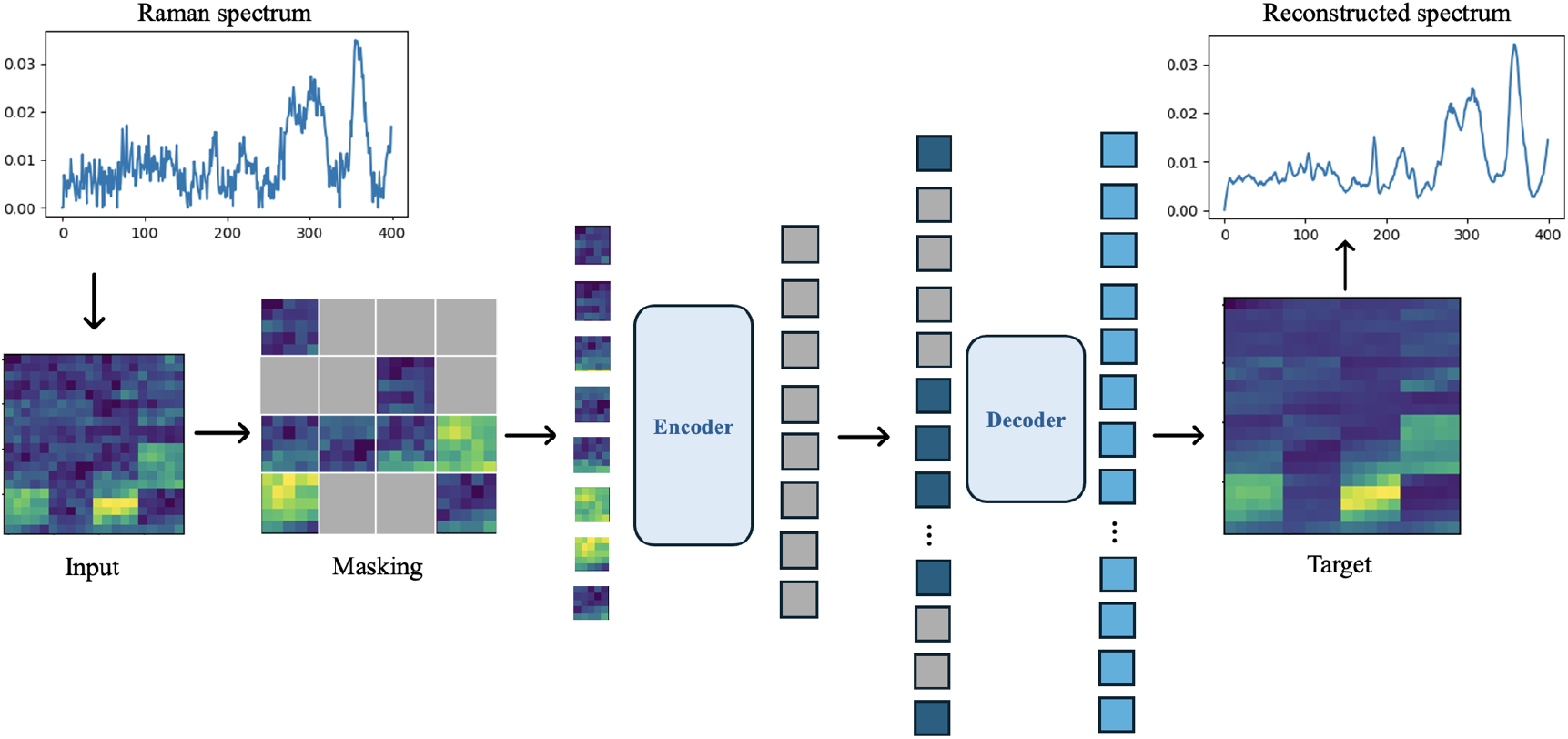
RamanMAE architecture is adapted from VIT-MAE with 1D spectra presented as 2D images obtained by transforming the spectra such that the contiguous 1D patches in the spectra spatially correspond to the 2D patches in the original VIT-MAE inputs.

## Methods

### Raman Datasets

We used published Raman datasets to train and evaluate the RamanMAE models in this study. The first dataset that we used was derived from the Horgan et al. DeepR publication (20). This dataset consisted of over 159k Raman spectra from multiple cells at high SNR and their matched low SNR counterparts acquired with smaller exposure. This dataset also has the corresponding Raman image coordinates, which we leveraged to visualize the embedding-based clustering outputs. We also employed the cartilage dataset used in the DeepR paper from a previous publication for demonstrating transferability (27). The second dataset we used to evaluate the relevance of the RamanMAE-derived embeddings for downstream training was derived from the publication by Paidi et al. (28). This dataset has single-cell Raman spectra from multiple cell lines of distinct metastatic abilities from a genetically matched model of breast cancer metastasis. This dataset has Raman spectra from distinct locations in parental MDA-MD-231 (P231) breast cancer cells, their circulating tumor cells (CTC) derived from the same tumor after implantation in mice, and lung metastatic cells (LM) that have successfully metastasized to lungs. This dataset consists of over 20k spectra from multiple cells and conditions. Third, we used two Raman spectral datasets derived from mouse tumors using a similar spectral acquisition protocol - (1) 16.5k spectra from mouse tumors of varying metastatic potential used in a prior publication, and (2) 5k spectra from mice treated with immunotherapeutic agent aPDL1 and corresponding controls (14, 15). We trained the RamanMAE on the former dataset and tested the transferability of the model for the prediction of therapy response in the latter dataset.

### RamanMAE architecture and training

Widespread adoption of emerging AI methods in life sciences requires turnkey solutions where possible. Therefore, in this study, we leveraged the standard VIT-MAE model (where code is publicly available) and made minimal changes to the architecture, such as input shape, layer size, and embedding dimensions, to train RamanMAE. To make the VIT-MAE, which takes 2D image data as input, compatible with 1D Raman datasets without significant architectural changes, we proposed a strategy to wrap the 1D spectra into a 2D image such that the contiguous 1D patches in the spectra spatially correspond to the raster scanned 2D patches in the original VIT-MAE architecture **Figure 1**. This setup ensures the equivalence with a 1D version of the VIT-MAE model, as the 2D image patches in VIT are flattened and positional encoded serially. We shaped 400-dimensional Raman spectra in the fingerprint wavenumber region to images of size 20×20 as inputs to our RamanMAE and reshaped the reconstructed decoder output images back to 400-dimensional Raman spectra. For each spectrum, we used a patch size of 16 in 1D spectra, which translated to a patch size of 4 in 2D image, yielding 16 patches for each spectrum. During the training of RamanMAE, we used a masking ratio of 0.5 to randomly mask half the region of the spectra in each step. To reduce the model complexity, we used an encoder projection dimension of 64, a decoder projection dimension of 32, 4 encoder heads, 3 encoder layers, 4 decoder heads, and 1 decoder layer. We used a batch size of 256, mean absolute error as the loss function, and AdamW optimizer to train RamanMAE. We trained the RamanMAE on a T4 GPU in Google Colab.

### Evaluation of RamanMAE

We trained RamanMAE using the VIT-MAE framework and custom functions in Tensorflow with Keras backend. We used the embeddings from the encoder output to perform the clustering of spectra into various groups using the k-means clustering method in the scikit-learn library in Python. The random forest classifiers were trained on the RamanMAE embeddings using classifiers in the scikit-learn library. For the leave-one-cell-out and leave-one-mouse-out classification, we iteratively train random forest models on the spectra from all the cells except one cell or mouse each time, which is left out and employed as a test set to predict the class label using the mode of all the spectral predictions. We used the UMAP library to generate the 2D UMAP projections from the RamanMAE embeddings.

## Results

### RamanMAE learns features of Raman spectra to enable accurate reconstruction of the masked patches and smoothing

We leveraged a large publicly available Raman spectral dataset consisting of pairs of high and low SNR spectra from biological cells to train and evaluate our RamanMAE model. We split the entire high SNR dataset into training, validation, and test datasets and trained RamanMAE using a masked token prediction objective while minimizing the loss of spectral reconstruction. Our training mean absolute error loss reduced asymptotically to 0.040 and the validation loss of 0.047. We tested the trained RamanMAE mode on an independent test dataset without masking the regions of spectra and observed excellent reconstruction of the spectra at the decoder output (**Figure 2A-F**). As expected, due to the denoising property of autoencoders, we also observed significant smoothing of the Raman spectra (PSNR = 27.88) while preserving important Raman spectral features of the cell spectra. Since the low throughput of Raman spectroscopy generally yields low SNR spectra without long acquisition times, we sought to test whether we could achieve similar performance when using the low SNR version of the spectra for reconstruction. We observed that the resultant smooth reconstructions were very similar to those obtained by performing the reconstruction on the high SNR spectra (**Figure 2D-G**). The high cosine similarity (0.98) between the reconstructed spectra from high and low SNR versions of the same spectrum further boosts our confidence in the RamanMAE reconstruction (**Figure 2G**). We compared the performance of RamanMAE-based smoothing with the Savitzky-Golay algorithm (which is sensitive to polynomial order and window size choices) and found that RamanMAE provides equivalent or slightly better similarity between reconstructions from the low and high SNR spectra (**Figure 2H**) (29). By varying the masking ratio from 0.25 to 0.75, we observed that the spectral reconstruction is not very sensitive to the masking ratio (**Figure 2I**).

**Figure 2.**
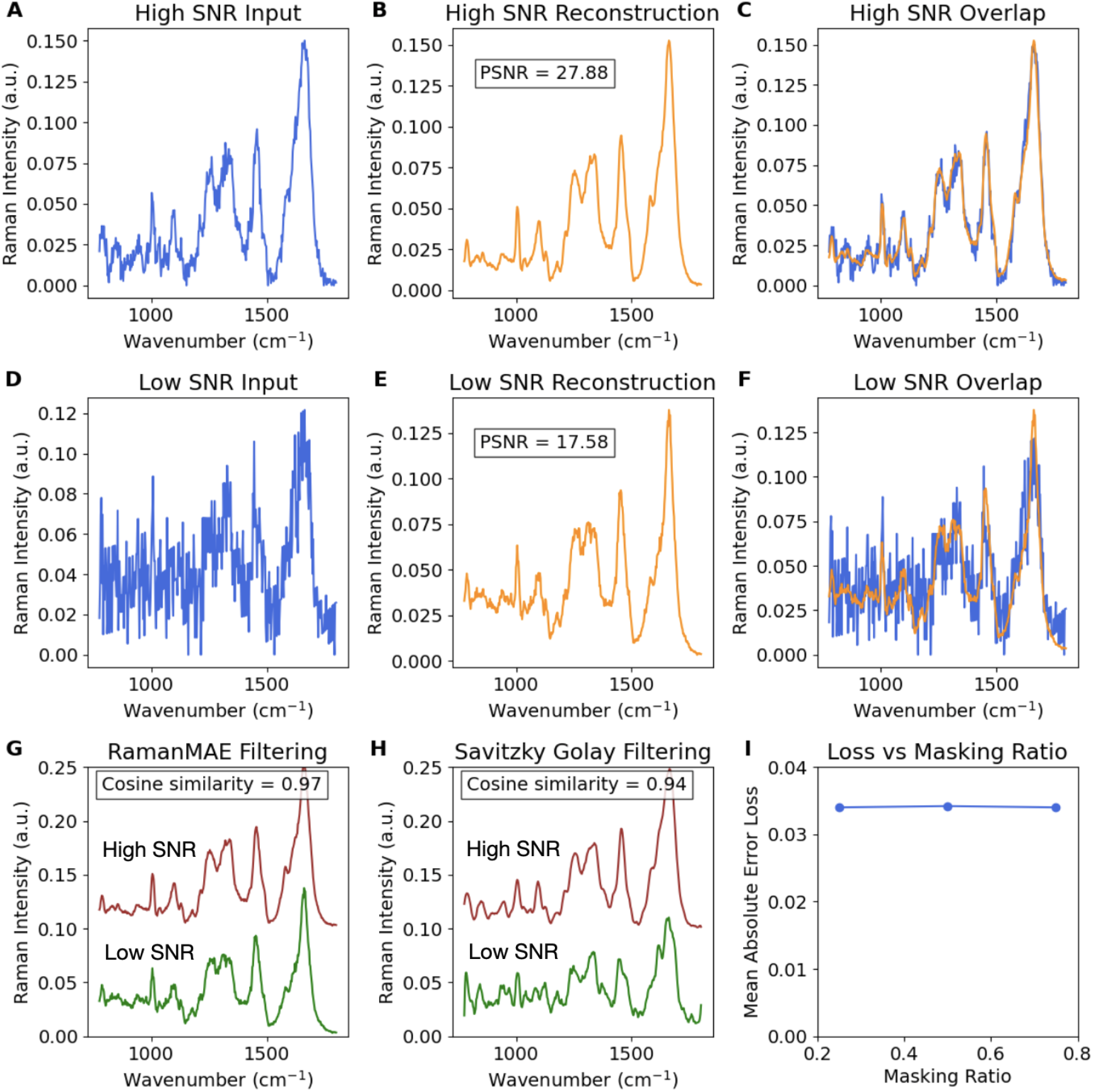
The representative examples of input spectra from the test dataset and their reconstruction at the Raman MAE decoder output are shown along with an overlay of the two for – (A-C) High SNR spectra, and (D-F) Low SNR spectra. (G) The RamanMAE reconstruction of high SNR and low SNR spectra is compared, and its cosine similarity is shown. (H) The Savitzky-Golay filtered (fifth order polynomial and 21 window length) versions of low SNR and high SNR spectra are shown for comparison. (I) The lack of significant impact of masking ratio on the test loss is shown.

### The learned RamanMAE embeddings capture features capable of identifying molecularly distinct regions of biological samples

The key advantage of masked autoencoders is their ability to robustly capture the domain-relevant embeddings of the underlying data at the encoder output. After successful training of RamanMAE, we inspected the embeddings obtained from the single-cell Raman spectra from the dataset. We performed k-means clustering in the low-dimensional embedding space and identified a clear demarcation between the spectra from the cells and the empty region in the hyperspectral dataset away from the cell (**Figure 3A**). We performed k-means clustering again on the identified cellular spectra with 3 components to evaluate if we could identify different compartments within the cell. Our reconstruction of the cellular image with the cluster labels reveals a clear correlation of the identified clusters with cellular components, specifically the nucleus, the cytoplasm, and the cell membrane that surrounds the cell.

**Figure 3.**
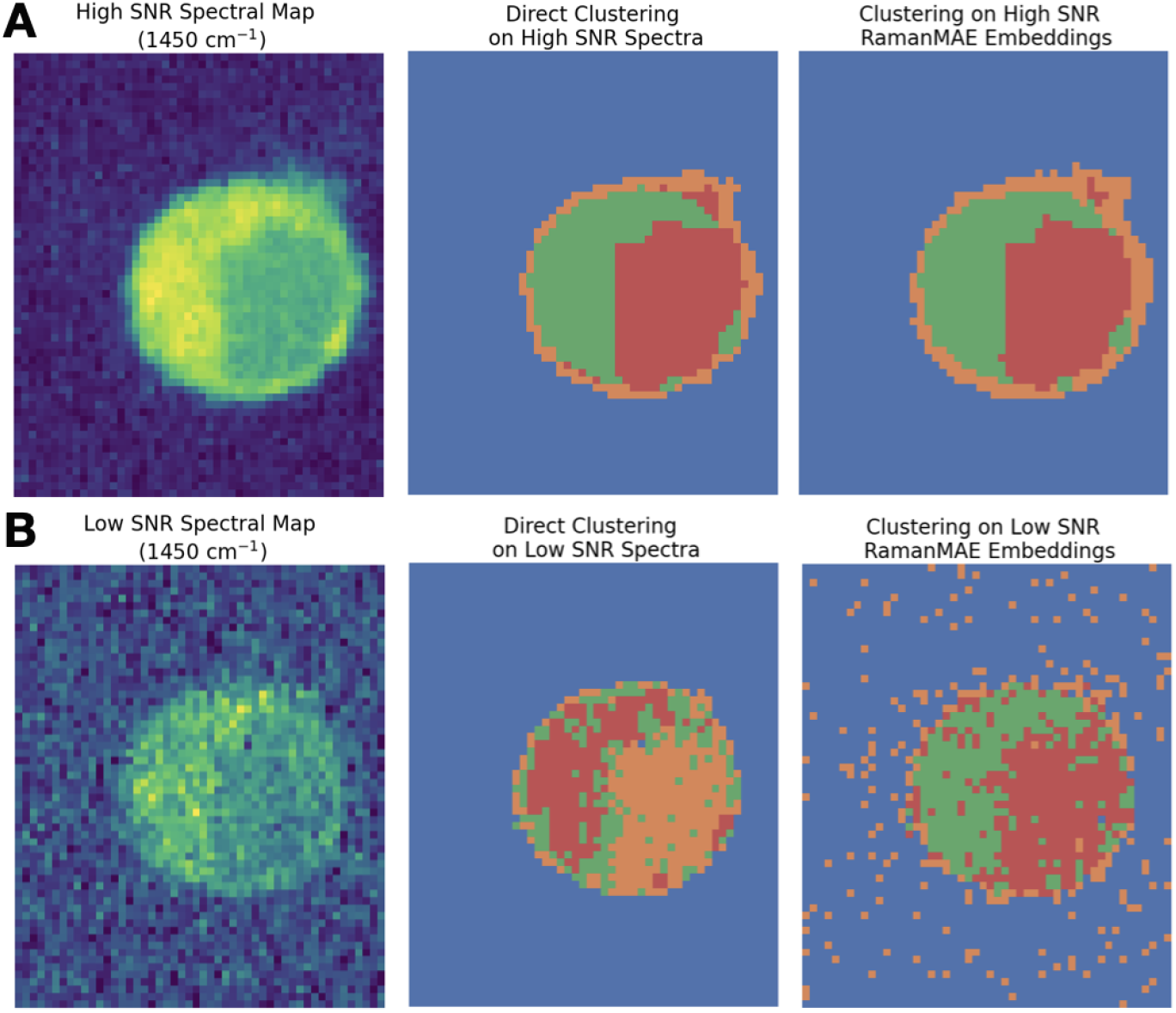
The spatial Raman image of a representative single cell is shown using univariate Raman peak intensity and cluster labels obtained from k-means clustering respectively, using full spectra and RamanMAE embeddings for – (A) High SNR data, and (B) Low SNR data.

Next, we used the corresponding low SNR spectra from the same cells to assess if the embeddings obtained from the low SNR spectra also provide similar clustering ability ((**Figure 3B**)). By using k-means clustering directly on the Raman spectra, we observed that the high SNR spectra were able to provide similar compartmentalization as RamanMAE, but the performance significantly deteriorated with the use of low SNR spectra, where the cell membrane doesn’t get as clearly demarcated and is misclassified with the nuclear compartment. We found that the RamanMAE embeddings obtained from the low SNR spectra provide similar compartmentalization as the high SNR spectra embeddings and enable meaningful demarcation of distinct cellular regions. These results indicate that RamanMAE learns biologically meaningful representations of the spectral data in addition to smoothing the data. We expect that the clustering algorithms in the embedding space can also leverage prior information about the underlying biology to choose hyper-parameters such as the number of clusters for specific biological applications.

### The learned RamanMAE embeddings allow accurate classification of Raman spectra

To evaluate the suitability of the RamanMAE embeddings for downstream classification tasks on challenging biological applications, we leveraged the published dataset comprised of Raman spectra from cancer cells of varying metastatic abilities. We used the decoder from the trained RamanMAE to extract the embeddings for the spectra belonging to multiple cells of parental MDA-MB-231 cells (P231 n=15), circulating tumor cells (CTC, n=15), and lung metastatic cells (LM, n=15). First, we use UMAP projection to reduce the dimensionality of the embeddings to 2D to visualize the clustering of the P231, CTC, and LM cells in the reduced dimensions. As expected, the UMAP projection shows that the P231, CTC, and LM cells are progressively clustered according to their metastatic potential and stage (**Figure 4A**). Since the circulating tumor cells will have a population of cells that will become lung metastatic cells in the future, a significant degree of overlap between CTC and LM cells is reasonably expected.

**Figure 4.**
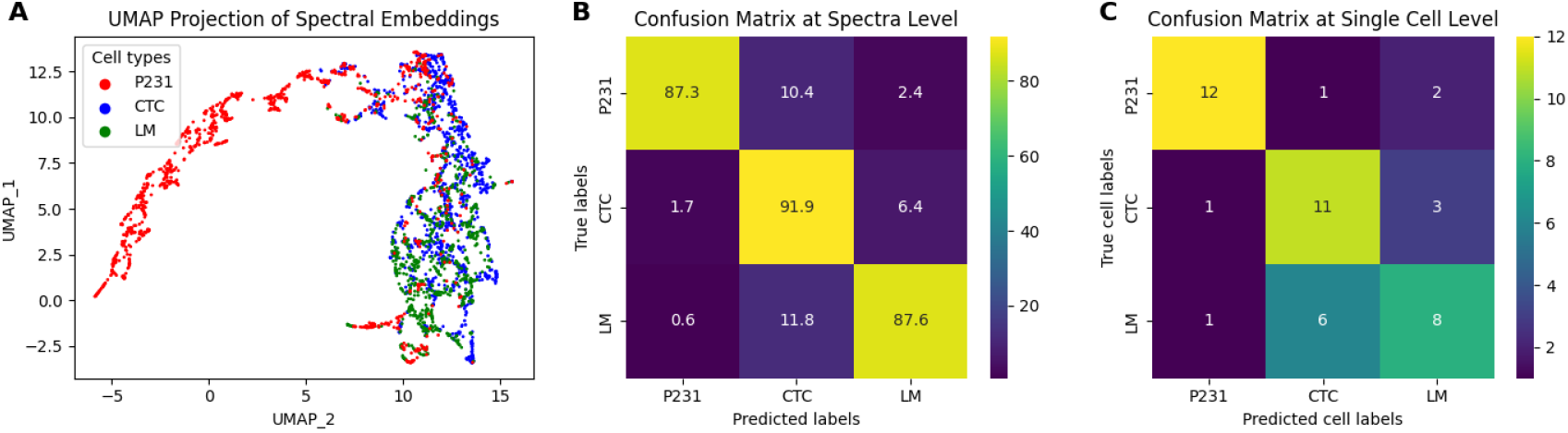
(A) The UMAP distribution of spectra in the RamanMAE embedding space is shown for the three classes. The confusion matrices for the random forest classification of cell types using the RamanMAE embeddings on a (B) per-spectrum and (C) per-cell basis are shown.

Next, we used random forest classification using the three class labels on a per spectra level and found that the classifier trained on the RamanMAE-derived embeddings can predict the spectra from the closely related cell types with high accuracy. The accuracy for the prediction of P231, CTC, and LM cell spectra were respectively 87.3%, 91.9%, and 87.6% on the independent test. The confusion matrix for the classification on a per spectra basis is provided in **Figure 4B**. Finally, since these spectral data are derived from single cells, not all the points within the cell of a specific type will have the class property, particularly in such closely related progressive classes, due to biological variability. Therefore, we trained the random forest classification on a per-cell level using a leave-one-cell-out approach, where we iteratively trained the random forest classification on spectra from all the cells but one cell that was used as a test cell for the derived classifier. For each left-out test cell, we provide the mode of the predictions as the predicted label. The confusion matrix from such leave-one-cell-out analysis shows high classification accuracy for P231 and CTC cell types and slightly poor performance for LM cells ((**Figure 4C**)). However, due to their similarity, the misclassified LM cells were consistently classified as the neighboring class CTC, which is consistent with the cluster overlap in the UMAP projection. Overall, these results show that the RamanMAE embeddings capture biologically relevant information and have utility for downstream classification tasks in challenging biological datasets.

### RamanMAE allows transferability of embeddings learned on an initial task to novel biological applications

We used the RamanMAE model trained on the high SNR cell spectra dataset to evaluate the biological relevance of embeddings of Raman spectra obtained from an articular cartilage tissue section in the same wavenumber region. The univariate Raman intensity maps derived from Raman wavenumbers representative of the nucleus, cytoplasm, and collagen are compared to the spatial distribution of RamanMAE embeddings (**Figure 5A-B**). We observe that the spatial distribution of some of the RamanMAE embeddings matches the known component maps very closely.

**Figure 5.**
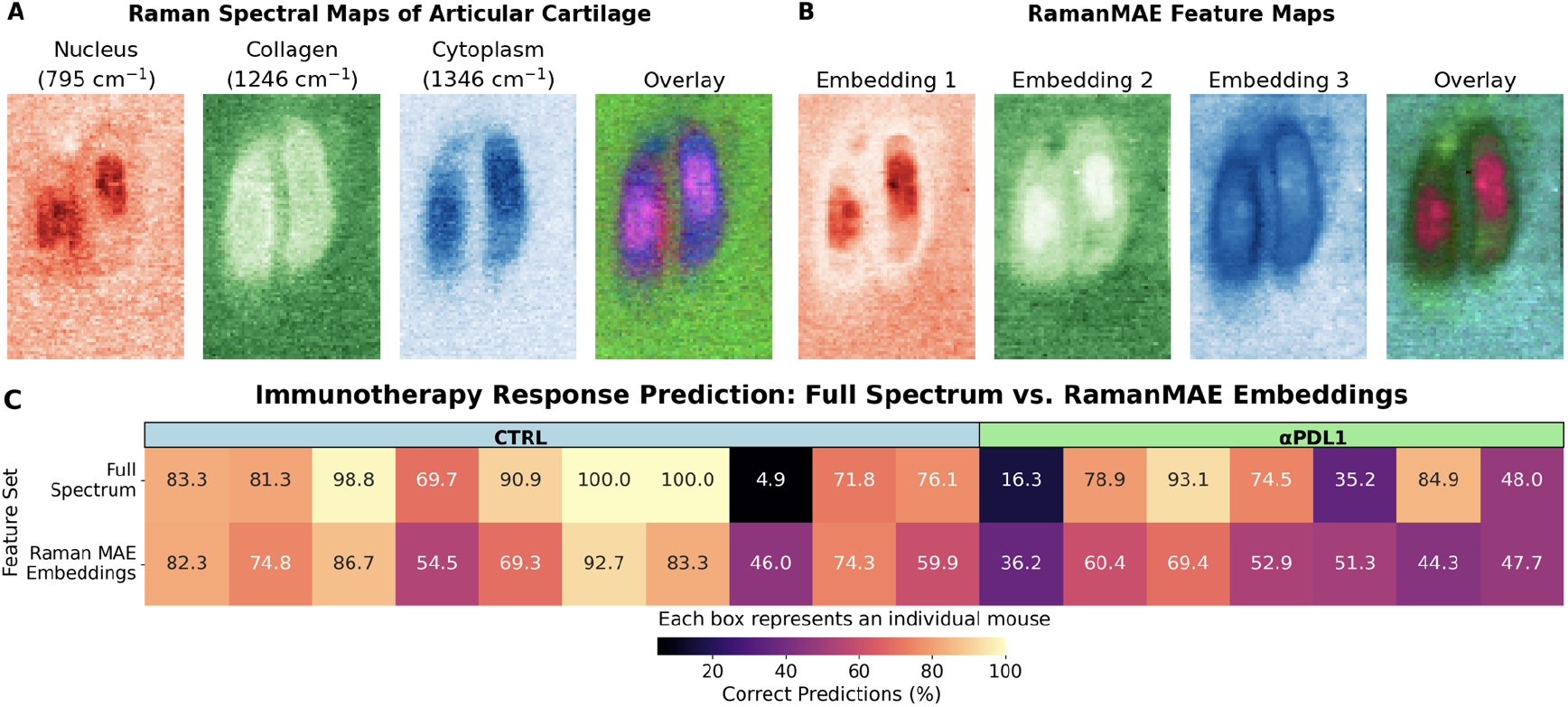
(A) Univariate Raman maps of the articular cartilage tissue at wavenumbers representative of the nucleus, collagen, cytoplasm, and their overlay are shown. (B) Feature maps of select RamanMAE embeddings from RamanMAE trained on cell data that resemble nucleus, collagen, and cytoplasm are shown along with their overlay. (C) The results of leave-one-mouse-out random forest analysis for predicting immunotherapy response for control mice (CTRL) and mice treated with *α*PDL1 are shown for both the full spectrum and their embeddings derived from the RamanMAE model trained on the metastasis dataset.

Next, we evaluated the transferability of the self-supervised RamanMAE model pre-trained on mouse Raman spectra from a biological study of cancer metastasis to a supervised classification task on a separate mouse study focused on therapy response prediction, where the Raman spectra were captured in the same wavenumber range. We trained random forest classifiers using a leave-one-mouse-out approach on both the original Raman spectra and their RamanMAE embeddings (obtained from the model trained on the cancer metastasis dataset) of mouse tumors treated with immunotherapeutic *α*PDL1 and their corresponding control (CTRL) mice. By comparing the percentage of correct predictions derived from subjecting Raman spectra and RamanMAE embeddings of the left-out independent test mice to the corresponding random forest models, we observed that the RamanMAE embeddings provide very similar levels of accuracy for predicting the immunotherapy response on a per-mouse level (**Figure 5C**). These examples illustrate the transferability of the RamanMAE model trained on one data-rich application to provide biologically meaningful embeddings of Raman spectra acquired at a different scale or in an unrelated biological application.

## Discussion and Conclusion

The success of advances in machine learning and artificial intelligence is tied to the widespread utility of the developed methods across a wide range of disciplines. The areas of chemical and biological sciences typically lag behind other areas in the adoption of new data-driven technologies, due to several factors such as a lack of large structured datasets, multimodality of experimental datasets, limited interpretability of complex models, and regulatory oversight in clinical applications (30, 31). Therefore, the tools and techniques developed in the field of machine learning must be robust to these factors for their successful deployment in such challenging environments. Therefore, in this paper, we propose the RamanMAE framework, which is adapted from VIT-MAE, for tackling challenges in the new domain of biomedical Raman spectroscopy.

We leveraged published large Raman datasets to demonstrate the feasibility of adapting masked autoencoders for the reconstruction of masked regions of Raman spectra and learning meaningful embeddings of the data. We showed that RamanMAE leverages the inherent denoising property of masked autoencoders to smooth the Raman spectra while preserving the subtle but sharp Raman features of the spectra. It is important to note that this denoising property is particularly relevant to Raman spectra due to the low throughput and resultant low SNR. We showed that the reconstruction efficiency is quite robust to noise in the Raman data, and the RamanMAE model trained on high SNR data is capable of obtaining sufficient smoothing of low SNR spectra, when subjected as a test set. Using a test dataset comprised of Raman spectra of cells, we confirmed that the learned embeddings capture biologically meaningful features of the Raman data, such as those that differentiate the compositional constituents of the cell membrane, nucleus, and cytoplasm. These results suggest that the models learned on high SNR spectra can be leveraged for denoising and clustering the spectra obtained at low SNR in future experiments, thereby saving the spectral acquisition time. Such time savings can allow us to obtain data from larger batches of cells, thereby more effectively capturing the biological heterogeneity and temporal dynamics at high resolution.

One of the biggest challenges in the field of Raman spectroscopy is the collection of large datasets for supervised classification, particularly due to the high dimensionality of the Raman spectra. Traditional dimensionality reduction methods like PCA and MCR-ALS are routinely employed to interpret the chemical and biological constituents of spectra prior to their use with machine learning methods (30, 32). Therefore, we next assessed the relevance of embeddings of data learned by RamanMAE for downstream applications due to the inherent dimensionality reduction by the encoder. We leveraged another challenging dataset of genetically matched cancer cells in different stages of metastasis and showed that the RamanMAE embeddings capture sufficient information to classify closely related cancer cell types. Furthermore, the 2D UMAP projections of the embeddings confirmed that the RamanMAE embeddings capture the continuity of the three classes in the dataset, consistent with their differences along the metastatic potential axis.

In this study, we also demonstrated that RamanMAE models built on large spectral datasets derived from one application can be utilized to perform dimensionality reduction of a spectral dataset from a different application if similar spectral acquisition parameters are chosen. Furthermore, by training the RamanMAE model on cell spectral data and using it to reconstruct the compositional map for a tissue dataset, we showed the transferability of the method across applications at different scales. Such transferability incentivizes researchers across labs and institutions to align on domain-specific spectral acquisition parameters and protocols to contribute to the creation of large and diverse shared datasets for training future spectral language models. The availability of such spectral language models will enable more applications with limited data (e.g., patient biopsy and surgical samples) to build machine learning solutions in the low-dimensional embedding space.

The field of Raman spectroscopy is currently undergoing a transformation due to the availability of larger datasets, accessible computing infrastructure to researchers, and demonstrated examples of success in extracting biochemically rich insights from spectral datasets that are of direct industry and clinical relevance. To demonstrate the ease of adoption of VIT-MAE for Raman spectroscopy, we built RamanMAE by making minimal changes to the VIT-MAE architecture. This approach will allow more researchers from chemistry and biology to adopt RamanMAE for their workflows without deep domain expertise in masked autoencoders. Although this paper did not focus on architectural enhancements for improving the properties of the method, we acknowledge that this turnkey method in its current form may not be the most optimized solution. Motivated by the success of RamanMAE, we plan to study the properties of this architecture further in future studies and explore the integration of multiple spectroscopic and microscopy modalities in the embedding space.

In conclusion, we demonstrated the relevance of the masked autoencoders for learning meaningful biological representations of Raman spectra. We showed that by making minimal changes to the VIT-MAE architecture, we can build a turn-key solution for Raman spectroscopy. We showed that the denoising property of the developed RamanMAE is useful to uncover subtle spectral features from low SNR spectra. We showed that the embeddings obtained at the encoder output capture sufficient chemical information for downstream clustering and classification tasks. We also demonstrated the transferability of the learned biological representations across scale and applications. Overall, this study shows the promise of masked autoencoders for learning a spectral language for biomedical applications.

## Data and Code Availability

The unpublished data and code will be made available via a public repository after the peer review and publication of the manuscript.

